# Insect Pollinators: The Time is Now for Identifying Species of Greatest Conservation Need

**DOI:** 10.1101/2023.10.20.563282

**Authors:** Phillip deMaynadier, Matthew D. Schlesinger, Spencer P. Hardy, Kent P. McFarland, Laura Saucier, Erin L. White, Tracy A. Zarrillo, Bruce E. Young

## Abstract

Severe declines in the abundance of insects, including economically and ecologically important pollinators are alarming conservationists and the public. Yet despite these increasingly well documented declines, relatively few pollinating insects other than some butterflies and bumble bees have appeared as Species of Greatest Conservation Need (SGCN) in State Wildlife Action Plans, decadal-scale blueprints for wildlife conservation efforts across the United States that require updating in 2025. Species absent from SGCN lists are ineligible for congressionally appropriated State Wildlife Grants that direct millions of dollars annually for their conservation. In the past, knowledge about the distribution and abundance of many insect pollinators was too poor to identify those meeting state guidelines for inclusion as SGCN. Using case studies from 4 northeastern states, we demonstrate that sufficient conservation status data now exists for many pollinators, including bees, butterflies, moths, beetles, and flower flies, to identify at-risk species meriting inclusion on SGCN lists in many states. Doing so will increase funding available for surveys, habitat protection and enhancement, and other conservation activities that will benefit this vitally important guild.

---

Alarming reports documenting insect declines, even in seemingly intact habitats, is a sobering new chapter in the global biodiversity crisis (Wagner 2020). An increasing body of evidence has led to a growing consensus that the declines are steep, taxonomically broad, and geographically widespread (Wagner 2020, Sanchez-Bayo and Wyckhuys 2021). Because insects perform so many functions in natural ecosystems, from pollination to decomposition to biological pest control, loss of insect abundance stands to have profound impacts on other organisms (Forister et al. 2019), including humankind. Researchers have suggested numerous ways to confront this crisis (Karahara et al. 2021). We argue that, in the United States, an important and underutilized conservation tool to protect insects are State Wildlife Action Plans (SWAPs). Broader representation of insects in SWAPs will hasten coordinated actions to protect them.

With the first plans submitted to the U.S. Fish and Wildlife Service in 2005, and requiring revisions every 10 years thereafter, SWAPs are a relatively new and important tool for preventing extinctions in the United States. SWAPs identify state vulnerable species and habitats and provide a blueprint for how state wildlife agencies mobilize and direct millions of dollars of conservation funds each year. A diverse coalition of government, tribal, academic, and private stakeholders typically contribute to SWAPs, resulting in consensus views on state wildlife conservation priorities. With such broad professional input, SWAPS are valuable tools for recovering Threatened and Endangered species listed under the U.S. Endangered Species Act and preventing declining species from needing to be listed under the Act.

Foundational to every SWAP is a list of Species of Greatest Conservation Need (SGCN) that are deemed most at risk in the state, requiring immediate management attention (USFWS 2020). States vary in their definitions of “wildlife” and therefore in the breadth of taxa included in SGCN lists. Vertebrates are always considered. While some freshwater invertebrates such as mollusks, crayfishes, and odonates are increasingly well covered (Mawdsley et al. 2017, Hamilton et al. in review), most insect groups are poorly represented in SGCN lists (Hamilton et al. in review). Although most states included some insect pollinators in their 2005 and 2015 SGCN lists, few states included more than a handful of these species, and important groups such as flower flies (Diptera: Syrphidae) were almost entirely absent (Table 1).

**Table 1.**
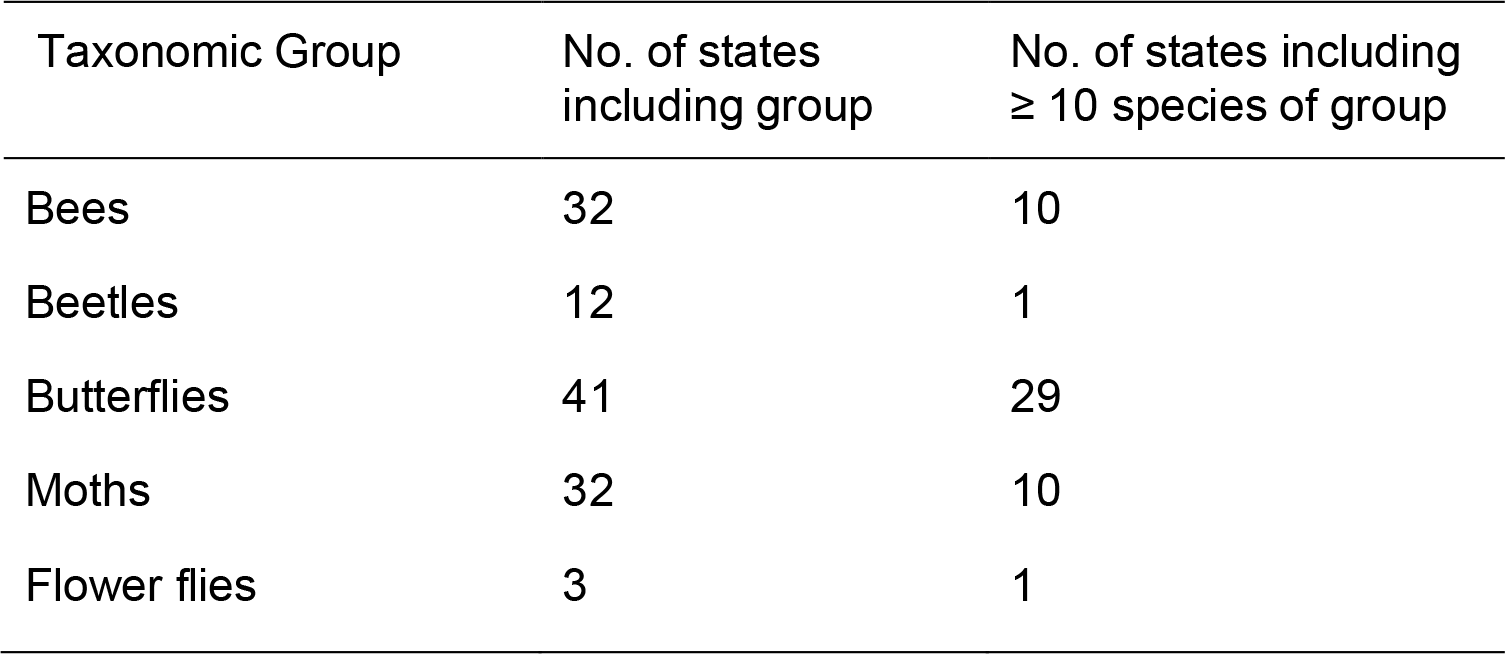
Representation of select insect pollinator groups in 2015 state lists of Species of Greatest Conservation Need (SGCN). Survey includes all 50 U.S. states. Beetles include only those known to be pollinators or belonging to families with known pollinators. Source: USGS (2021).

We argue here that reasons for excluding insects from previous SGCN lists, such as a lack of information or a popular premise that their needs are largely covered by conserving habitat for wide-ranging vertebrates, can now be discounted in many cases. The boom in reliably identified records available through online community science (also known as “citizen science”) platforms, digitized museum collections, as well as professional, targeted field surveys have greatly increased the knowledge base available for determining which species are rare or most at risk (Young et al. 2019). In addition, guidance for how to include pollinators in SWAPs already exists (The Heinz Center 2013). We use examples of pollinating insects from 4 states in the northeastern U.S. to illustrate how knowledge about the status of insect populations is rapidly expanding. Pollinators perhaps receive more research attention than other insect groups due to their economic importance (Gallai et al. 2009, Jordan et al. 2021), but as a guild they include both well and poorly known species similar to other insect guilds. And because this growth in knowledge about pollinators is not unique to the Northeast, we discuss the urgent need for SWAP committees nationwide to include greater coverage of at-risk pollinators in their review of SGCN lists in forthcoming (2025) State Wildlife Action Plan revisions.

## Advances in knowledge about northeastern U.S. pollinators: Case studies

As is true globally, insects and other invertebrates dominate the biota of North America, generally exceeding 95% of the animal species diversity in most regions (Stein et al. 2000). Furthermore, in many U.S. states the legal definition of “wildlife” includes insects, presenting a daunting challenge to capacity and expertise for government authorities. As a result, state wildlife agencies often triage their limited resources toward the conservation of invertebrates by focusing on those taxa that: a) are relatively well-studied, b) include multiple imperiled species, and c) are known to provide critical ecosystem services. Pollinating insects are a flagship example of one such group. To illustrate recent advances in knowledge about the distribution and status of insect pollinators, we share case study investigations from 4 states in the northeastern U.S. While states vary in taxonomic foci and approaches to gathering information, the end result is similar: greater understanding of the state’s diversity of insect pollinators and identification of species that are likely to be experiencing population declines and/or be in danger of becoming locally extirpated.

### Maine

Butterflies are among the few insect groups that have benefited from considerable attention by Maine naturalists, starting with collections from as early as 1870 and recently via community science contributions during the Maine Butterfly Survey (MBS) completed in 2016. Data generated from the effort originated from 3 primary sources: a) community scientists, b) targeted, professional surveys, and c) curated material from private collections and museums. The goals of the MBS were two-fold, to raise public awareness and concern for butterflies and pollinating insects in general, and to increase scientific knowledge of the state’s butterflies, thereby improving efforts to protect them. Key outcomes included an annotated checklist of Maine’s 120 species, updated NatureServe subnational conservation status ranks (S-ranks; Faber-Langendoen et al. 2012) and state legal status, a specimen collection curated by the Maine State Museum, and a database of all known Maine butterfly observations (>34,000 records) (Figure 1). Many of these records are noteworthy, including >240 new county records, 12 new state records, and 1 new U.S. national record (Short-tailed Swallowtail, *Papilio brevicauda*). Of special interest is the relatively high proportion (∼20%) of butterflies considered Endangered, Threatened, Special Concern, or Extirpated, most of which are also SGCN in Maine’s Wildlife Action Plan. Finally, a published atlas was recently produced summarizing the biogeography and conservation concerns of the region’s fauna entitled “Butterflies of Maine and the Canadian Maritime Provinces” (deMaynadier et al. 2023).

**Figure 1.**
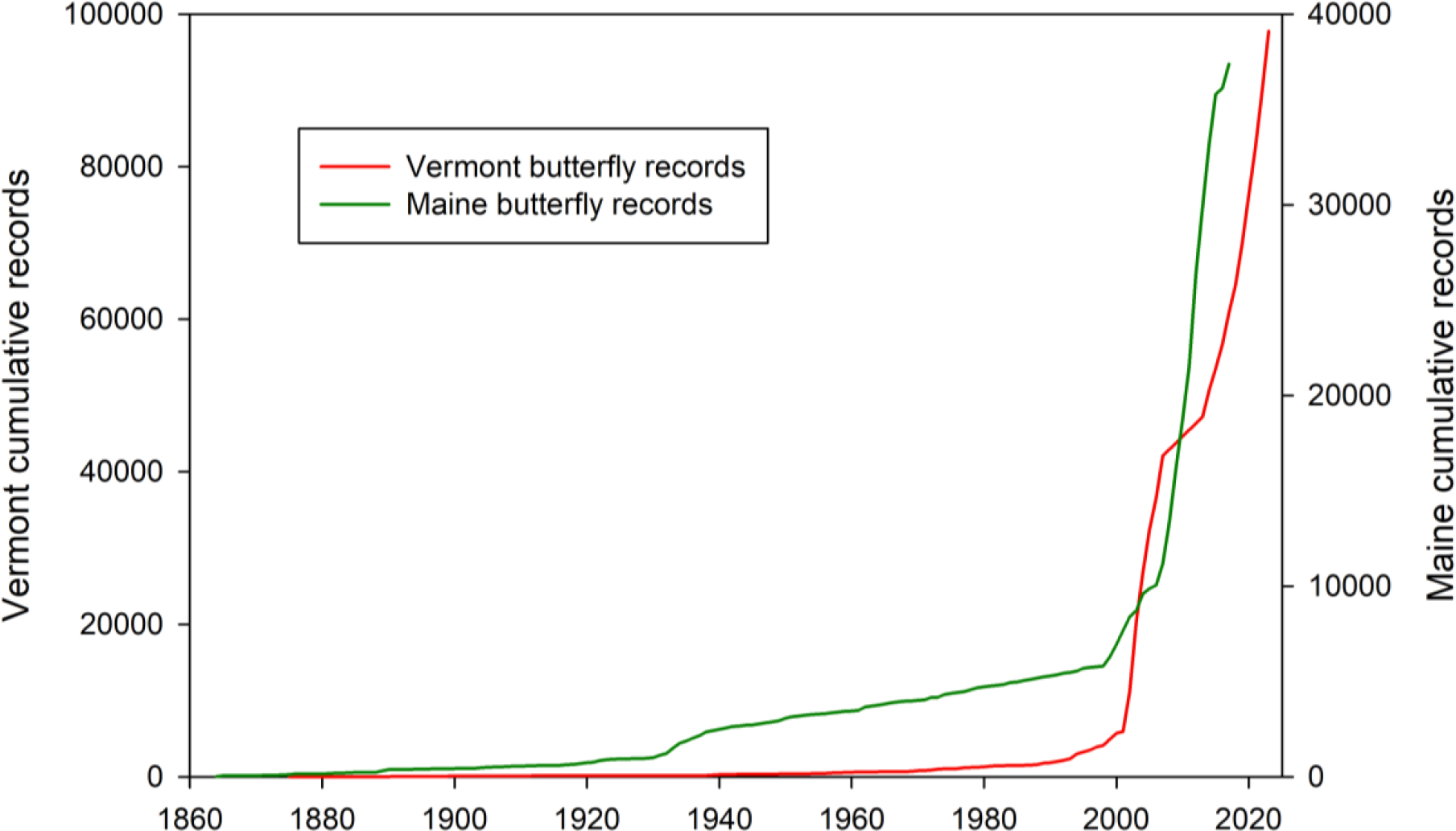
Cumulative number of butterfly records in Maine and Vermont from 1860 to 2021, indicating the recent dramatic increases associated with community science atlasing projects in both states.

Maine is fortunate to have a relatively good understanding of another important pollinator group -- bees. Using data compiled from publications, museum collections, and field surveys, a recently published county checklist reports 278 species from across 6 families (Dibble et al. 2017). Much of the research attention afforded bees in Maine is motivated by their role as pollinators, especially of Lowbush Blueberry (*Vaccinium angustifolium*), an economically important crop. More recently, the Maine Department of Inland Fisheries and Wildlife (MDIFW) has focused its attention on assessing the state’s bumble bee fauna (*Bombus* spp.). A recently completed statewide community science survey -- the Maine Bumble Bee Atlas (MBBA) -- significantly advanced our knowledge of the group’s distribution and status in the state. In 2015, MDIFW partnered with the University of Maine to coordinate volunteers in the collection of bumble bee specimens and photo vouchers. From 2015-2020, over 300 participants contributed more than 27,000 new records, confirming that 15 of the 17 species historically recorded in Maine are extant. The American Bumble Bee (*B. pensylvanicus*) and federally Endangered Rusty Patched Bumble Bee (*B. affinis*) are considered potentially extirpated. Combining MBBA data with global conservation status information from NatureServe, Maine has assigned SGCN status to 10 of its bumble bees. As is the case across North America (Goulson et al. 2015), it is thought that habitat loss, pesticides, introduced diseases and parasites, and intensive agriculture have played a role in bumble bee declines in Maine.

Most recently, MDIFW is investigating its flower fly fauna (Family Syrphidae), an especially important pollinator group in the Northeast (Klymko et al. 2023). In support of this effort, NatureServe contracted with the Atlantic Canada Conservation Data Centre (ACCDC) to develop a checklist of flower flies in Maine using records gleaned from the literature, museum collections, online community science platforms (e.g., iNaturalist, BugGuide), and recent surveys. As a result, Maine now has a database of >3,500 records of flower flies comprising 214 species, each with newly assigned S-ranks. While most of Maine’s Syrphids do not have enough records or life history information to develop informed ranks (NatureServe rank SU; see https://explorer.natureserve.org/AboutTheData/DataTypes/ConservationStatusCategories for NatureServe conservation status rank definitions), 8 species were flagged as species of conservation concern (ranked as SH, SX, S1, and S1S3). With this new information, MDIFW is prepared to consider the addition of at-risk flower flies to the state’s SGCN list during the 2025 SWAP update, thereby leveraging State Wildlife Grant funds for increased research and conservation of a previously under-studied taxon.

### Connecticut

Connecticut has the good fortune to have been home to numerous prolific Lepidopterists over the years, many associated with Yale University. Consequently, there is a relatively impressive knowledge base of Connecticut’s butterfly fauna, with some records dating back to the late nineteenth century. Capitalizing on a growing interest in butterflies by the public, the Connecticut Butterfly Atlas was conducted from 1995 through 1999 with the purpose of better understanding the distribution and relative abundance of butterflies statewide. One of the first such atlases to be published in the U.S. (O’Donnell et al. 2007), it focused on documenting changes in the distributions of butterflies at the state level utilizing historical occurrences to compare with atlas-related survey efforts. Survey design was based on both the Massachusetts Audubon butterfly survey (https://www.massaudubon.org/nature-wildlife/insects-arachnids/butterfly-atlas) and the Atlas of Breeding Birds of Connecticut (Bevier 1994). One hundred thirteen species were documented pre-atlas and 110 species documented during the project; noteworthy are species likely lost from the state (regal fritillary [*Speyeria idalia*], silvery checkerspot [*Chlosyne nycteis*]), species that experienced geographic range shifts (common ringlet [*Coenonympha tullia*], Ocola skipper [*Panoquina ocola*]) and non-native species establishment (European skipper [*Thymelicus lineola*]) (O’Donnell et al. 2007). The Connecticut Butterfly Atlas will serve as a baseline for assessing future faunal shifts due to global climate change and continued anthropogenic fragmentation of habitats.

Connecticut also has a rich historical baseline of its bee species community, dating back to the mid-19th century. Early publications include species descriptions of bees from Connecticut by renowned natural historians such as E. T. Cresson, W. H. Patton, C. Robertson, and T. D. A. Cockerell. In 1916, the first checklist of the bees of Connecticut was written by Viereck et al. (1916), including information on species distribution by county, known floral records and phenology, and additional species expected to occur in the state. Intensified interest in Connecticut’s modern day bee fauna arose following the unsettling report from the National Research Council (2007) regarding the uncertain status of pollinators in North America. Shortly thereafter, scientists at the University of Connecticut (UCONN) (funded through State Wildlife Grants) began conducting bee surveys in habitats such as sand barrens and early successional land under utility corridors. Five bee species were added to Connecticut’s Endangered, Threatened and Special Concern List in 2010 as a direct result of these early survey efforts. Notably, Connecticut was the first state in the U.S. to include bees on its state Endangered species list. Additionally, six bee species were included as SGCN in the 2015 Connecticut SWAP— *Bombus ashtoni* (= *bohemicus*), *B. affinis, B. pensylvanicus, B. terricola, Macropis ciliata*, and *Epeoloides pilosulus*. In response to the state listings, a long-term wild bee monitoring program was initiated in 2010 by the Connecticut Agricultural Experiment Station (CAES) and continued until 2021. This program was based on a standardized protocol designed for the U.S. Fish and Wildlife Service (USFWS) and included cooperators at 6 locations in representative habitats across the state. Additionally, surveys of coastal beaches and marshes, bogs and fens, agricultural fields, planted pollinator meadows, natural grasslands, and forests were also conducted by researchers at CAES, the Connecticut Department of Energy and Environmental Protection, Yale University, and UCONN. The data obtained from these surveys, as well as a thorough literature review (see Zarrillo et al. 2023 for references), records from private collections, observation records from online community science websites such as iNaturalist, and the digitization of major museum collections (Ascher 2016, Gall 2023, CAES 2023) — CAES, UCONN, the American Museum of Natural History, the Peabody Museum of Natural History at Yale University, and the Museum of Comparative Zoology at Harvard University — informed the recent documentation of Connecticut bees (Zarrillo et al., in review) and the assignment of state conservation ranks using NatureServe protocols.

Of the approximately 384 bee species documented for Connecticut, at least 43 species have not been detected in the state since 2000 (Zarrillo et al., in review). Of the 124 focal bee species reviewed for conservation assessment, 22 species were given a conservation rank of S1, 7 species are considered likely extirpated (*B. affinis, B. ashtoni* [= *bohemicus*], *Hylaeus basalis, H. saniculae, Coelioxys funerarius, Dianthidium simile*, and *Holcopasites illinoiensis*), 36 species are considered historical, and 13 species (all pollen specialists) were designated SU. These conservation ranks can help identify at-risk bees for potential designation as SGCN, and the subsequent allocation of resources for further surveys, monitoring, and habitat restoration efforts.

### New York

New York has long been a focus for entomological studies, facilitated by the presence of two major collections—the Cornell University Insect Collection and the American Museum of Natural History—that house tens of thousands of New York specimens dating back to the mid-1800s. Many of these specimen records have been digitized (e.g., Ascher 2016) and periodic faunal checklists (e.g., Leonard 1928) have yielded a good baseline for evaluating changes in species presence and distribution for a variety of pollinator taxa.

Attention to native pollinator conservation accelerated in New York upon the inclusion of several species of bumble bees on the state’s SGCN list in 2015, inspired by Schweitzer et al.’s (2012) report on the status of North American species and subnational conservation status ranking by the New York Natural Heritage Program (NYNHP). In its 2016 Pollinator Protection Plan (NYS DEC and AGM 2016), the state called for a survey of native pollinators and funded the NYNHP to design and implement it. The 5-year Empire State Native Pollinator Survey (White et al. 2022) used hundreds of field surveys, thousands of community science observations, and extensive compilations of data from museum collections and partners to inform the conservation status (S-ranks) of 457 pollinator species representing bees, flies, beetles, and moths. The project built on previously published research (e.g., Leonard 1928, Matteson et al. 2008, Bried and Dillon 2012, Ascher et al. 2014, Young et al. 2017), and digitized museum collections (e.g., Ascher 2016) to map distributions of focal taxa.

Approximately 60% of the focal species assessed were found to be at risk of extirpation from New York, with bees and flies comprising the most imperiled groups. Several new species were added to state lists, but 25 species of bees, 23 flies, 22 beetles, and 9 moths were considered historical (NatureServe status rank SH) because of a lack of post-1999 records. For 59 species, data were too scarce to be assigned a conservation status rank. These S-ranks should help inform the designation of many new SGCN insects during the upcoming revision of New York’s SWAP. Additional products included distribution maps for each species showing current (post-1999) records compared to historical (pre-2000) county records plus simple phenology charts for when adults were documented to be active in the 2 time periods. Requests for data have been frequent and report downloads have been in the hundreds, highlighting the increasing interest from natural resource agencies, scientists, and the general public for information on pollinating insects.

### Vermont

For centuries, humans have celebrated and documented the diversity of life that helps define Vermont. Over the last ∼50 years, more than 25,000 Vermonters—professional biologists, amateur naturalists, students, and others—have contributed to detailed statewide wildlife surveys and atlases. Collectively they have amassed over nine million primary biodiversity records, one of the most critical components for informing regional conservation rankings and policies.

Most statewide pollinator surveys and atlases have largely been conducted under the umbrella of the Vermont Center for Ecostudies’ Vermont Atlas of Life project (VAL), in partnership with the Vermont Fish and Wildlife Department and others. All VAL projects share data through the Global Biodiversity Information Facility (GBIF), allowing data to be incorporated into a wide range of additional studies.

The diversity and conservation status of butterflies were largely a mystery in Vermont before hundreds of volunteer naturalists joined the first Vermont Butterfly Atlas (2002-2007; Figure 1) (McFarland and Zahendra 2010). Atlas-generated occurrence data facilitated the designation of 16 butterflies as SGCN for the Vermont SWAP (Kart et al. 2005, VTFWD 2015). Butterfly observations shared after the atlas with crowd-sourced platforms like e-Butterfly and iNaturalist continued to provide important information, including 4 new species for the state. Twenty years later, the Second Vermont Butterfly Survey (2023-2027) is poised to detect changes in the fauna’s distribution and abundance. Already, the second atlas has discovered a new species for Vermont, Bog Elfin (*Callophrys lanoraieensis*), likely to be added to the state’s list of SGCN.

The Vermont Bumble Bee Atlas (2012 - 2014) digitized thousands of specimens from historical collections and conducted standardized surveys to assess the current status and distribution of the 17 bumble bee species known from the state. The results of this work (Richardson et al. 2019) documented the recent extirpation of 3 species and declines in several others, leading to the designation of 9 bumble bees as SGCN for the 2015 Vermont SWAP (VTFWD 2015). VAL recently expanded this work to the entire bee fauna and launched the Vermont Wild Bee Survey (2019 - 2022). This project combined standardized surveys by volunteers as well as targeted surveys by professional biologists and resulted in the first comprehensive checklist for the state’s bee fauna (Hardy et al., unpublished data). Dozens of new state records, including a globally rare (G2) species - *Megachile rugifrons* were first documented by the Vermont Wild Bee Survey. Using records from the survey and approximately 65,000 bee records available through GBIF, VAL generated subnational conservation status ranks for 350 species, which helped to inform a preliminary list of 55 bees to designate as SGCN for the 2025 revision of the Vermont SWAP (Hardy et al 2022).

## Status of northeastern pollinators

Northeastern states’ efforts to inventory and assess the conservation status of pollinators have led to a growing scientific basis for selecting pollinating insect species that meet state criteria for designation as SGCN (Figure 2). Variation in funding availability and programmatic priorities has led to heterogeneity in the depth at which different taxa have been addressed to date. All of the states profiled have begun assessing bees (Hymenoptera), ranging from bumble bees only (Maine) to multiple bee groups (New York, Vermont, and Connecticut). Strikingly, high numbers of the bees examined are at risk of extirpation. All 4 states have also reviewed the status of dozens of butterflies and moths (Lepidoptera). Two states, Maine and New York, have assessed flower flies (Diptera: Syrphidae), with Maine assessing the fauna comprehensively and New York focusing on forest-dependent species. Although most Maine flower flies did not have sufficient information to calculate a conservation status rank, 6 species are now considered at risk and under consideration for SGCN status. New York’s statewide survey resulted in the identification of many species at risk. Additionally, only 1 of the profiled states (New York) has assessed the conservation status of flower-visiting beetles (Coleoptera; Cerambycidae; Subfamily: Lepturinae and Scarabaeidae), finding many species to be at risk.

**Figure 2.**
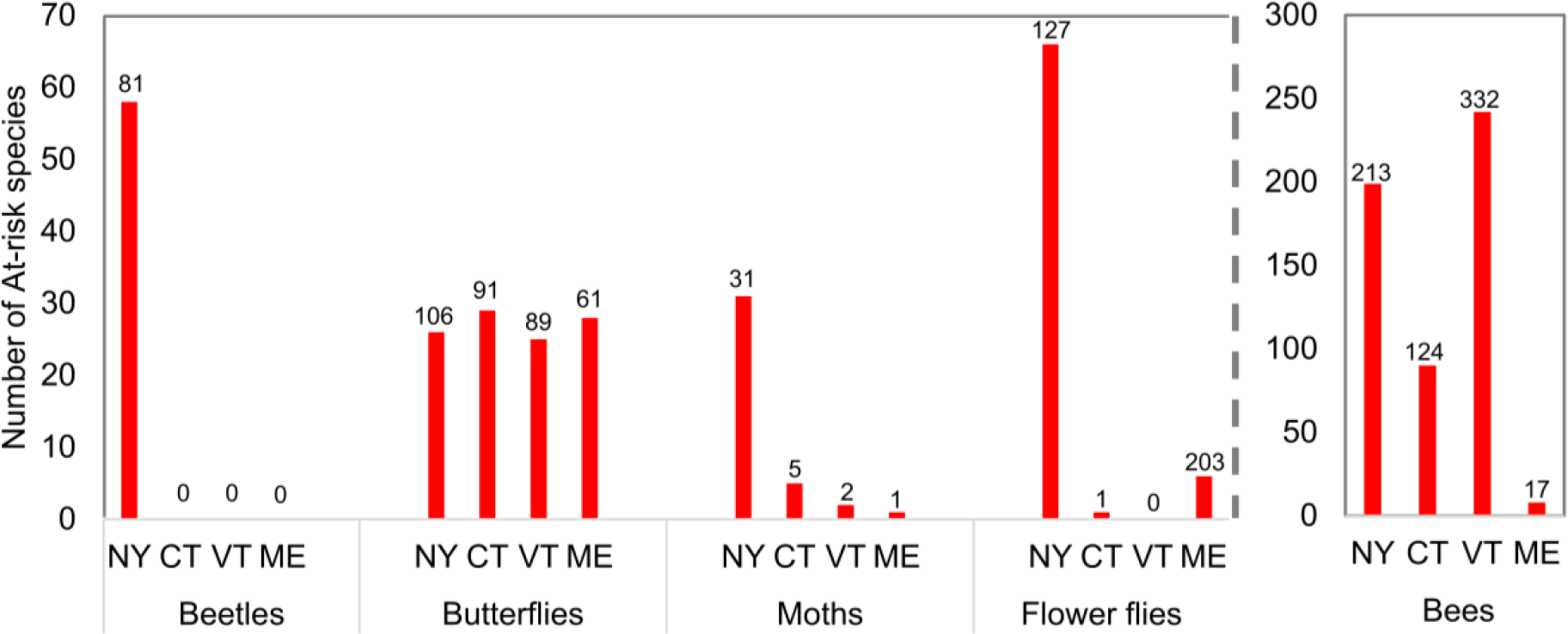
Progress on assessing state conservation status of major insect pollinator groups in selected northeastern U.S. states as measured by the number of species assessed as at-risk of extirpation. Column numbers refer to the total number of species in each group assessed. Definitions: At risk = NatureServe subnational conservation status rank of SH, S1, S2, or S3. The flower fly number for New York includes 10 bombyliid flies (3 of which are at risk) in addition to flower flies. Beetles include flower longhorn (Lepturinae) and selected scarabs (genus *Trichiotinus*) only. Most of the species in these groups serve as pollinators.

## Insect pollinators as Species of Greatest Conservation Need

Although each state determines rules for inclusion in its SGCN list, criteria in most states include documentation of rarity, steep population decline, or significant life history vulnerabilities (e.g., habitat specialization). An assessment of at-risk status can occur at various geographic scales, for example at the level of the state (e.g., state-listed as Endangered, Threatened, or Special Concern), region (e.g., U.S. Endangered Species Act; western butterfly assessment [Forister et al. 2023]), or world (e.g., NatureServe global conservation status or IUCN Red List of Threatened Species category; Faber-Langendoen et al. 2012, IUCN 2012). Our case studies demonstrate a rapidly growing understanding of the distribution and status of insect pollinators in the northeastern U.S. Similar growth in pollinator knowledge is taking place across North America (National Research Council 2007) – knowledge that can be used to help justify greater inclusion of pollinating insects in SWAPs, alongside conservation recommendations for their recovery (The Heinz Center 2013). For species that have not yet been assessed by the methods above, agency biologists can identify new species that meet state SGCN criteria directly or by working with a state natural heritage program or other conservation partners (e.g., Universities, conservation nonprofits, focal taxon experts) to facilitate assessments.

The existence of a NatureServe subnational conservation status rank, a measure of the likelihood of a species becoming extirpated in a state, is one indication that enough information might be available to determine whether a species could qualify as a SGCN. State natural heritage programs, which are typically embedded in state government agencies to provide information on rare species, use the NatureServe conservation status methodology (Faber-Langendoen et al. 2012) to assign such ranks. Across the U.S., as of 23 June 2023, fully 1,718 unique pollinating species (bees, butterflies, moths, flower flies, and pollinating beetle groups) had 7,095 subnational status ranks assigned (a species can have as many subnational ranks as the number of states it occurs in; ranks of “SU” and “SNR,” for “unrankable” and “not ranked,” respectively, excluded). All 50 states and the District of Columbia have subnational status ranks for at least some pollinating insects (Table S1, available in Supporting Information). Although only a fraction of all U.S. insect pollinators have been evaluated, this species total nevertheless indicates that a substantial number of species may be well enough known for considering SGCN designation, within and beyond the 4 states profiled. Designating these species as SGCN would make SWG funding available for further inventory of key pollinator groups, which will improve knowledge of species’ distributions and potentially reveal new populations of at-risk species.

Key to effective state-level conservation planning and actions are the interconnections between a strong knowledge base for species and their habitats, status assessments leading to SGCN designation, and associated funding through State Wildlife Grants and other sources that enable work towards these assessments, and their recovery (Figure 3). The relationship among these various components of conservation can be illustrated by considering bumble bees in Vermont. For example, many bumblebee species in North America declined dramatically in the 1990s (Cameron et al. 2011). But lack of detailed quantitative data on historical and modern abundance and distributions in northeastern states hampered status assessments that were needed for conservation plans and actions. In Vermont, a concerted effort was made prior to the 2015 SWAP to fill this void with the launch of the Vermont Bumble Bee Atlas (Richardson et al. 2019). Biologists and trained community scientists then conducted standardized surveys across the state to assess their current status and distribution. The results of this work documented the recent extirpation of 3 species and declines in many others, leading to the designation of 9 of the state’s 17 species as SGCN for the 2015 Vermont SWAP (VTFWD 2015) and listing 4 species as state Threatened or Endangered, heralding a new era for their conservation. As a result of these designations, public and private land management plans began to take bumble bee habitat into consideration, new pesticide regulations and bans were enacted, and a statewide population monitoring program was launched. Following Vermont’s lead, many other states have or are currently atlasing their bumble bee fauna, including a 14-state effort initiated by Xerces Society with 8 more states embarking on atlas projects in 2024 (Xerces Society 2023). These atlases can inform and inspire states to take the first steps toward recovery planning and conservation actions for a heretofore understudied group.

**Figure 3.**
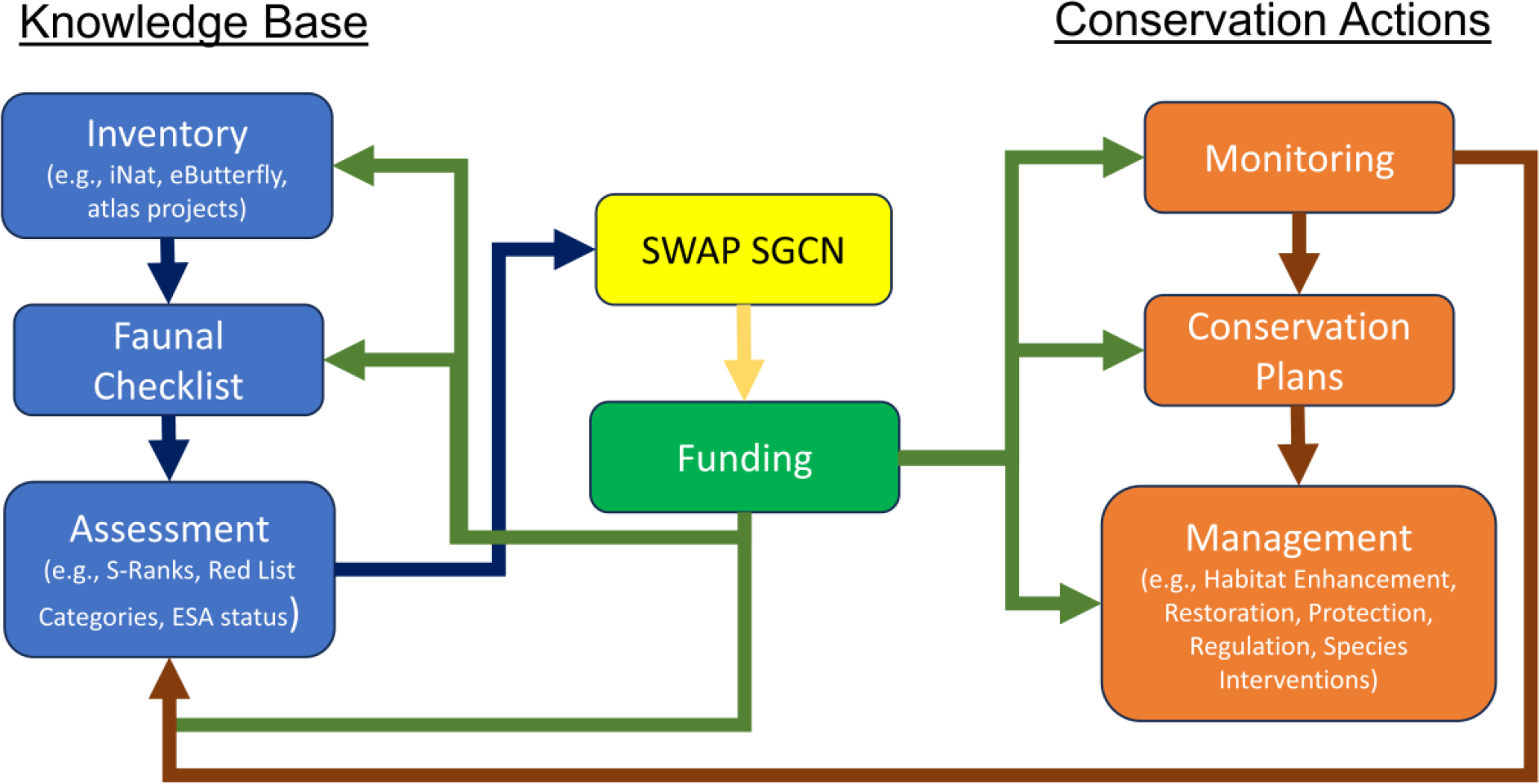
Relationships among knowledge, SGCN designation in State Wildlife Action Plans, funding (via State Wildlife Grants and other sources), and conservation actions by U.S. states. Abbreviations: S-Ranks = NatureServe subnational conservation status ranks; ESA = U.S. Endangered Species Act.

The conservation benefits of including pollinators and other insects as SGCN are many (Figure 3). First, shining the spotlight on these groups will help remind the conservation community of their ecological importance and vulnerability. Second, as mentioned, species designated SGCN are eligible for congressionally appropriated State Wildlife Grants. This funding totals approximately $60 million annually (USFWS 2020) and serves as the primary source of support for nongame and Threatened and Endangered species work in most state wildlife agencies. State Wildlife Grant monies can be used for further surveys, threat assessments, monitoring, habitat protection and management, and other measures that help support the recovery of at-risk species. Third, publicizing the fragile conservation status of a species by placing it on an SGCN list can catalyze research and management efforts on these and related species by partner institutions such as tribes, universities, and non-governmental organizations. In addition, federal agencies with multiple use mandates, such as the Bureau of Land Management and Forest Service, often consider the needs of state priority species as expressed in SGCN lists in their land-use and management decisions.

Collectively, these efforts constitute a broad conservation community response to the challenges presented by increasingly well-documented insect declines. Concern for pollinating insects is widespread and growing, having expanded beyond the scientific community to include farmers, politicians, journalists, and the general public. With states on the cusp of instituting once-in-a-decade revisions to their SWAPs, now is the time to convert scientific knowledge into action by designating a greater number of pollinating insects as SGCN.

## ACKNOWLEDGMENTS

Funding was provided by a Sarah K. de Coizart Article TENTH Perpetual Charitable Trust grant to NatureServe and State Wildlife Grants to Connecticut, Maine, and Vermont administered by the U.S. Fish & Wildlife Service. We thank Margaret Ormes for help with NatureServe’s biodiversity databases, Ron Butler for assistance with Figure 1, and Sarina Jepsen for reviewing an earlier draft of the manuscript.

